# A long non-coding RNA in the *let-7* complex acting as a potent and specific death effector of cancer cells

**DOI:** 10.1101/2021.07.16.452600

**Authors:** Tosca Birbaumer, Tommy Beat Schlumpf, Makiko Seimiya, Yanrui Jiang, Renato Paro

## Abstract

The *let-7* complex in *Drosophila* encodes three evolutionarily conserved microRNAs: *miR-100, let-7*, and *miR-125*. These act as heterochronic genes in regulating developmental timing in response to the steroid hormone ecdysone and play important roles in cell differentiation. Here we identify two additional long non-coding RNAs in the *let-7* complex, we named *let-A* and *let-B*. Both are transcribed in the large first intron of the primary RNA encoding the microRNAs. We show these RNAs to be sequentially expressed in early pupal stages in response to ecdysone signaling, albeit exhibiting a different expression pattern compared to the microRNA *let-7*. Surprisingly, ectopic expression of *let-A* in *Drosophila* cancer cells induces rapid cell death. Dead cells further release RNA molecules in the medium that is becoming toxic to other cancer cells. *In vivo* grown tumors lose their tumorigenicity after being incubated in the *let-A* induced medium. Moreover, feeding flies carrying transplanted tumor cells with such induced medium leads to reduced growth of tumors in a subset of hosts. Our results uncover a new lncRNA which can act as a potent and specific cell death effector for *Drosophila* tumor cells.

## Introduction

*lethal-7* (*let-7*) encodes an evolutionarily highly conserved microRNA (miRNA) and was among the first miRNAs to be discovered (Reinhart et al., 2000). In *Drosophila, let-7* is expressed from the let-7 complex (*let-7-C*), which is a single locus with a primary RNA encoding two additional miRNAs: *miR-100* and *miR-125* (Bussing et al., 2008). At metamorphosis, expression of these miRNAs is activated by the steroid hormone 20-hydroxy-ecdysone (ecdysone) (Sempere et al., 2002). Studies in *C. elegans* and *Drosophila* have shown that *let-*7 is required for controlling developmental timing and for cell differentiation (Bussing et al., 2008). Flies containing a complete deletion of all three miRNAs are viable and appear morphologically normal. Mutant adult flies, however, display behavioral and aging defects. The miRNAs expressed from *let-7-C* were found to be required for neuronal development, neuromuscular remodeling, and adult lifespan and aging (Chawla et al., 2016; Gendron & Pletcher, 2017; Sokol et al., 2008). In mammals, all three miRNAs are jointly present as large gene families scattered on different chromosomes in the genome. Due to the functional redundancy and complexity, the exact expression pattern and the biological relevance of the mammalian homologues of all three miRNAs remain unresolved (Roush & Slack, 2008). However, mis-expression of *let-7* or *miR-125* have been observed in several human cancers, indicating a possible function of these miRNAs in controlling cell proliferation and differentiation (Roush & Slack, 2008).

*Drosophila* is a powerful model to study the mechanisms underlying tumor formation and development (Gonzalez, 2013; Miles et al., 2011; Read, 2011). The central brain of *Drosophila* has been used to study neural stem cell-derived brain tumors in *brain tumor* (*brat*) gene mutants (Bello et al., 2006; Betschinger et al., 2006; Lee et al., 2006). The larval imaginal discs are also widely used to study epithelial tumors. For instance, cells mutant for the *polyhomeotic* (*ph*) gene, a member of the *Drosophila* Polycomb group, overproliferate in the larval eye-antennal discs and give rise to neoplastic tumors after transplantation (Classen et al., 2009; Martinez et al., 2009). We showed that *ph* loss of function mutant (*ph*^*505*^) cells in the developing larval eye-antennal discs are unable to differentiate and are trapped in an immature embryonic-like condition (Torres et al., 2018). Surprisingly, we discovered that an intrinsic tumor eviction mechanism marks the tumorigenic, undifferentiated cells for destruction (Jiang et al., 2018). At metamorphosis, ecdysone-induced expression of *the let-7-complex* transforms tumorigenic *ph*^*505*^ mutant cells into non-tumorigenic cells which become subsequently eliminated during the adult stage. A critical part in the process plays by the miRNA *let-7* which down-regulates among others the gene *chronologically inappropriate morphogenesis* (*chinmo*), an important driver of tumorigenesis (Narbonne-Reveau et al., 2016).

Large-scale sequencing and mapping of a number of different genomes uncovered extensive regions of transcriptions (Djebali et al., 2012). While many of these RNAs could be assigned to already well-known functional families, like mRNAs, tRNAs, snoRNAs or miRNAs, the majority of these transcripts remain functionally uncharacterized (Derrien et al., 2012; Jia et al., 2010). A particular class, long non-coding RNAs (lncRNAs), shows many hallmarks of mRNAs; cap-structure and polyA tail or the usage of the same transcriptional and post-transcriptional machinery, except for the capacity to encode the information for a protein (Ulitsky & Bartel, 2013). There is increasing evidence, however, that lncRNAs also fulfill a broad range of cellular and organismal functions as well (Kopp & Mendell, 2018; Liu et al., 2017). While in the past the conservation of the open reading frame in mRNAs often supported the functional characterization of the associated proteins across species, the much-reduced evolutionary preservation of lncRNA sequences hampers this task substantially. Often the function is restricted to a characteristic secondary structure allowing the interactions with specific partner molecules (Carlevaro-Fita et al., 2020; Hezroni et al., 2015). Hence, complex research strategies are often required to reveal underlying functional roles of specific lncRNAs (Kopp & Mendell, 2018).

In this report, we identify two new long non-coding transcripts in the *Drosophila let-7-C*. The two lncRNAs, which we named *let-A* and *let-B*, are encoded in the first intron of the *let-7-C* locus. We show that these lncRNAs are transiently expressed during normal development in response to ecdysone, but the expression patterns of both lncRNA have different dynamics to that of the miRNA *let-7*. Interestingly, induced expression of one of the two RNAs, *let-A*, results in rapid death of *in vitro* cultured *Drosophila ph*^*505*^ cancer cells, but not in S2 and Cl8 cells. Dead cells further release RNA molecules into the medium that then becomes toxic to other cancer cells. *In vivo* grown tumor tissue becomes non-tumorigenic after *in vitro* incubation with *let-A* conditioned medium. In addition, feeding such induced medium to host flies carrying transplanted tumors leads to reduced tumor growth.

## Results

### The *let-7* complex encodes two putative novel non-coding RNAs

The *let-7* complex in *Drosophila* is located on the second chromosome. The primary transcript encoding three miRNAs (*let-7, mir-125*, and *mir-100*) spans ~17 kb of genomic region and consists of three exons and two introns (Fig. 1A). In our previous study, we demonstrated that the *let-7-C* locus plays an important role in an endogenous tumor eviction mechanism (Jiang et al., 2018). Ecdysone signaling during metamorphosis, activating transcription of the locus and in consequence expression of the miRNAs, was found to be central for the process of transforming tumorigenic *ph*^*505*^ cells into non-tumorigenic “metamorphed” *ph*^*505*^ cells. At the adult stage, metamorphed cells eventually become eliminated by programmed cell death. We collected eye-antennal imaginal discs from early pupae as well as GFP-marked metamorphed cells from young adults and subjected them to a detailed transcriptome analysis by RNA-seq. To our surprise, we observed many reads mapping to the intronic region of *let-7-C* primary transcript (Fig. 1A). These reads were particularly enriched in the 5’ region of the first intron in one group of samples, and in part of the second half of the first intron in another (Fig. 1A). This indicates that the *let-7-C* may encode, besides the pri-RNA for the miRNAs, additional non-coding transcripts. In our previous studies, we showed that larval *ph*^*505*^ mutant cells can continue to proliferate and give rise to neoplastic tumors after being transplanted into adult host flies (Jiang et al., 2018; Torres et al., 2018). For further examination, we established a *ph*^*505*^ mutant cell line from this tissue (see material and methods). To test if ecdysone can induce the expression of the non-coding transcripts observed in the metamorphed cells, we treated *ph*^*505*^ cell culture with ecdysone overnight, isolated the polyadenylated RNAs, and performed transcriptome analysis by RNA-seq. This analysis showed many reads mapping to the second half of the first intron (Fig. 1A, ph^505^ 16h), suggesting ecdysone signaling can induce the expression of certain non-coding transcripts. Furthermore, an analysis of modENCODE data for RNA Pol II ChIP of *Drosophila* adult heads and Kc167 cells reveal two regions with strong Pol II accumulation within the first intron (Fig. 1A). These observations suggest that the *let-7-C* locus is more complex and encodes at least two lncRNA transcripts in addition to the primary transcript for the three miRNAs (Fig. 1A). Based on 5’- and 3’mapping results, as well as sequencing reads, we named one lncRNA *let-A* which consists of the first exon and the first half of the intron, and the other transcript *let-B* which consists of sequences in the second half of the first intron (Fig. 1B). Interestingly, three previously discovered ecdysone responsive elements (EcRE), which are required for the ecdysone-induced expression of *let-7-C*, are all located inside the *let-B* transcription unit (Fig. 1B).

**Figure 1.**
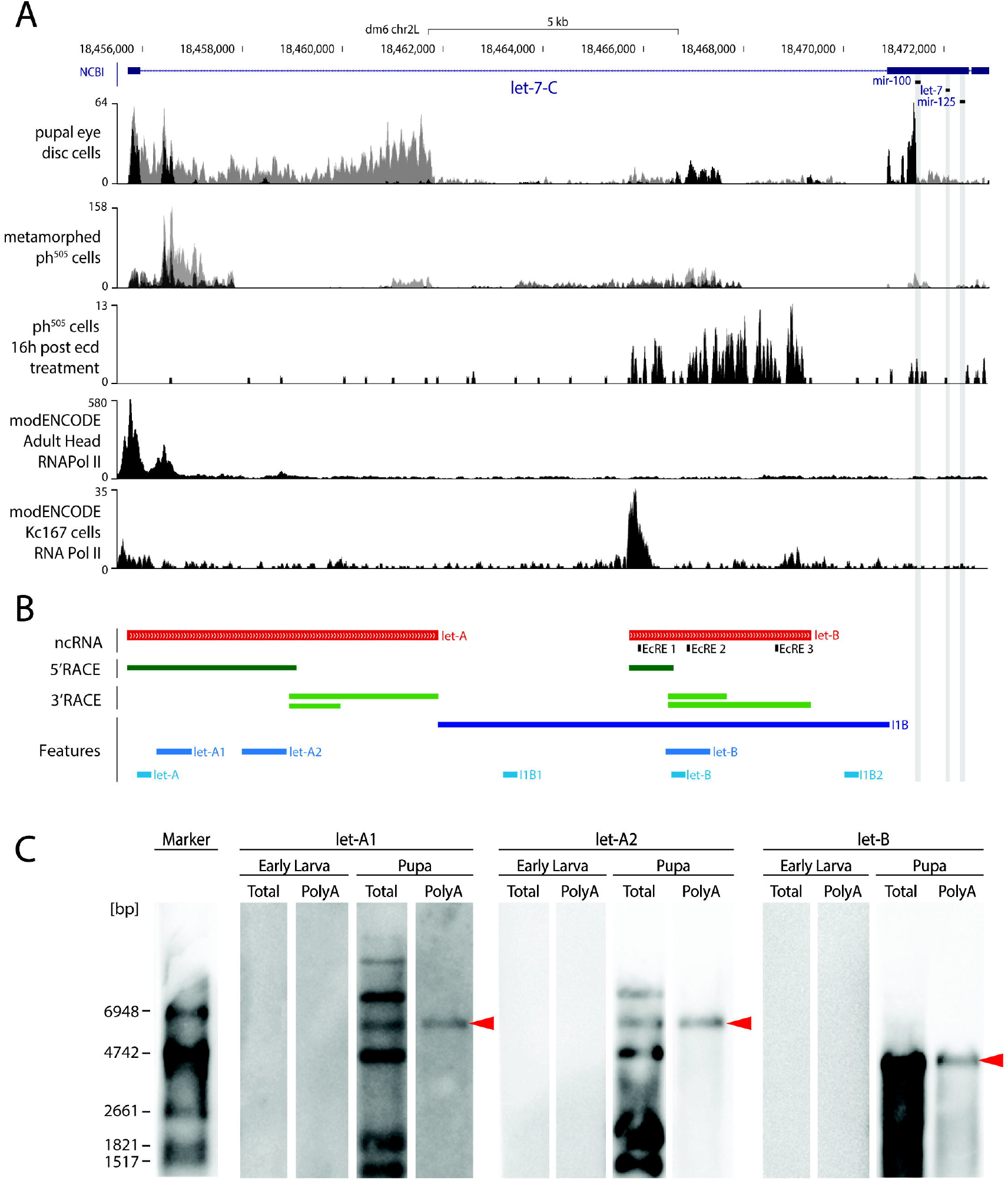
The *let-7-Complex* encodes two undocumented non-coding transcripts. (**A**) Genomic region of the *let-7-C* in *Drosophila*, containing 3 exons (blue boxes), one large intron and one small intron (dashed line). The three miRNAs (*mir-100, let-7*, and *mir-125*) are encoded in the second exon. Transcriptome analysis of wild-type eye-antennal disc cells at 48 hours after pupal formation (pupal eye disc cells) and metamorphed *ph*^*505*^ cells by RNA-seq revealed many reads (black and grey bars indicate two replicates) mapping to the first intron. RNA-seq analysis of ecdysone-treated *ph*^*505*^ culture cells (*ph*^*505*^ 16h post ecd) also showed sequencing reads mapping in the first intron. Analysis of modENCODE data for RNA Pol II ChIP of adult heads and Kc167 cells reveals multiple regions with strong Pol II localization within the first intron, indicating potential transcription start sites. (**B**) Based on the Pol II localization sites, the sequencing reads, and the 5’ and 3’ mapping experiments we named one lncRNA *let-A* and the other transcript *let-B* (red bar). Note the transcript *let-B* contain three known ecdysone responsive elements (EcRE1-3). 5’ RACE (dark green) and 3’ RACE (light green) revealed the 5’ end and 3’ end of both lncRNA transcripts. The partial intron sequence (dark blue), from the 3’ end of let-A to the 3’ end of the intron 1, is names as I1B and used as control in experiments described below. In features, let-A1, let-A2, and let-B are probes used for Northern analysis. let-A, I1B1, letB, and I1B2 are regions for qPCR analysis in Figure 2. (**C**) Northern analysis using the probes indicated in B. *let-A* and *let-B* are enriched in pupal polyA+ RNA (red arrow). The additional bands in the total RNA lanes suggest possible processing products of *let-A*, lacking the polyA-tail.

To confirm the *in vivo* existence of the two RNAs, we purified total as well as poly-A^+^ RNA from wild-type larvae and early pupae and performed Northern blots. Using two different probes for *let-A* (Fig. 1B), a transcript of about 6 kb in both the total and poly-A RNA samples was revealed (Fig. 1C). The transcript was detected only in the pupal stage but not in the larval stage. Longer exposures reveal also the smaller transcript isoforms (supplementary figure A). Interestingly, a number of additional bands are observed in the total RNA pool. This could suggest a further processing of the transcript into isoforms lacking a poly-A^+^ tail. Conversely, a probe for *let-B* revealed a single transcript of around 4 kb at the pupal stage (Fig. 1C). Hence, Northern results confirm the expression of the two lncRNA during normal development.

In order to determine the sequence of the two RNA transcripts, we performed rapid amplification of cDNA ends (RACE) analyses for both *let-A* and l*et-B*. Due to the composition of the *let-7-C* intron sequences, we were unable to sequence the *let-A* or *let-B* transcripts in their entirety, and only able to acquire overlapping fragments with either 5’ end or 3’ end (Fig. 1B).

### Expression profile of the two lncRNAs during development

To determine if these new non-annotated transcripts can be found in other transcriptome database, we searched publicly available RNA-seq data in Flybase, which had been generated from various tissues and cells at different developmental stages. Indeed, the intronic sequences can also be observed in a few samples and cell lines (Fig. 2A; see also Fear, JM and Oliver, B., GEO accession: GSE117217). Interestingly, the intronic sequences were detected particularly in samples from pupal stages, but less in larval and not in embryonic or adult stages, correlating again the expression of these transcripts to ecdysone signaling at metamorphosis.

**Figure 2.**
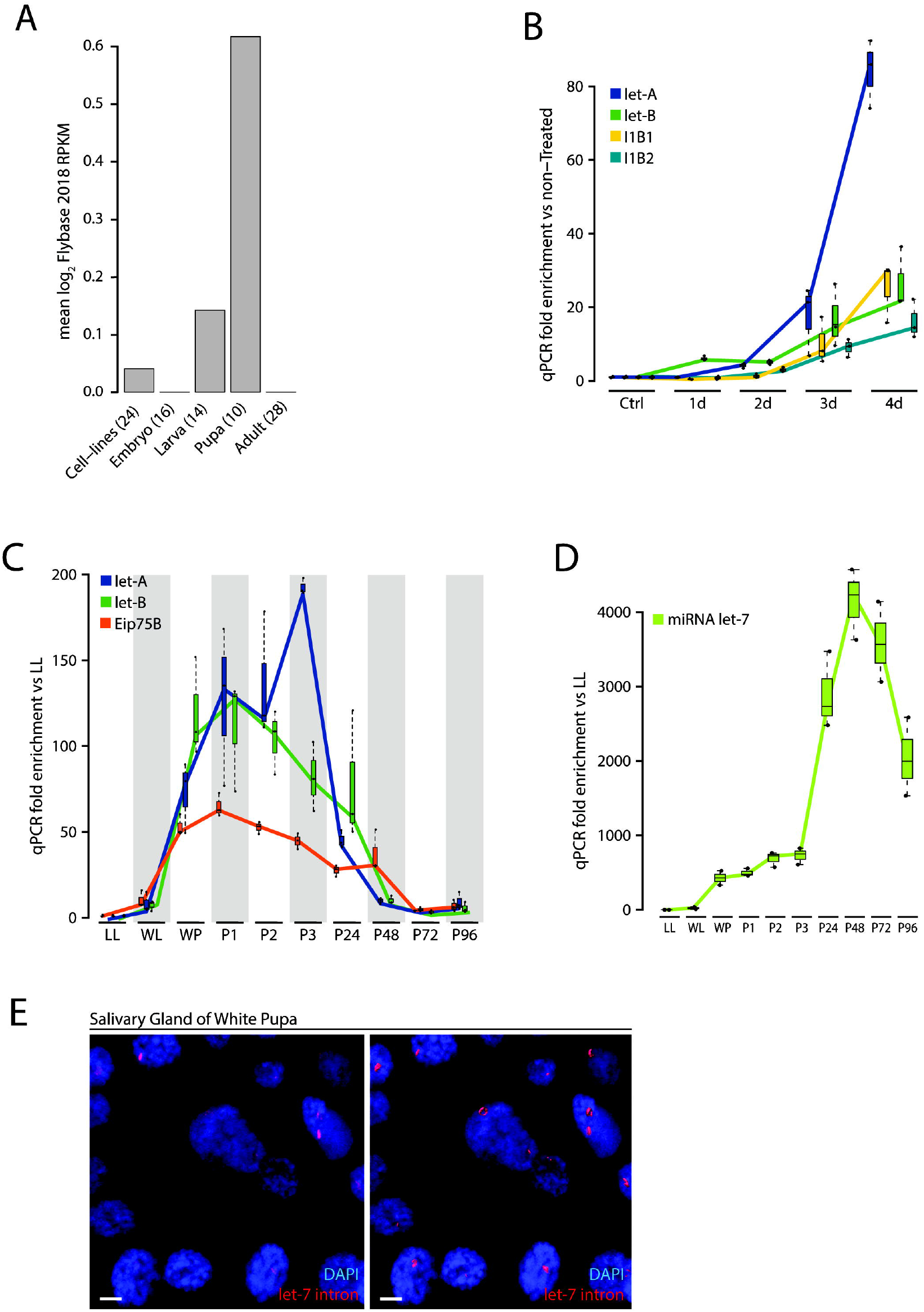
Dynamic expression of *let-7-C* lncRNAs during development. (**A**) *let-7-C* intron sequences can be found in publicly available RNA-seq data in Flybase. Notably, the intron sequences are enriched in samples from pupal stages, but not in embryonic, larval, and adult stages. (**B**) qPCR quantification of the expression levels of different intronic regions of the *let-7-C* in ecdysone-treated *ph*^*505*^ culture cells at different time points (Ctrl: untreated; 1d: 1 day; 2d: 2 days; 3d: 3 days; 4d: 4 days). Primers were used to amplify *let-A* coding region (purple), *let-B* coding region (green), intermediate region between *let-A* and *let-B* (I1B1, yellow), and 3’ region outside of *let-B* (I1B2, blue). qPCR fold enrichment was normalized to the untreated. (**C**) qPCR quantification of the expression levels of different intronic regions of the *let-7-C* in whole larvae or pupae from different developmental stages. Primers used were the same as in (**B**). qPCR fold enrichment was normalized to the expression at LL. *Eip75B*, a known ecdysone-induced gene, was used as a reference. LL: late 3^rd^ instar larvae; WL: wandering larvae; WP: white pupae; P1, P2, …: corresponding hours after pupae formation. (**D**) Expression dynamic of *let-7* miRNA at different developmental time points. (**E**) Confocal image of *in situ* hybridization of intron 1 sequences to whole salivary glands of white prepupae shows a strong signal (red) at the periphery of the nuclei (DAPI, blue). Right panel: 3D reconstruction of the same image using Imaris. Scale bar is 5 μm.

To further examine if the two lncRNAs become activated upon ecdysone signaling, we treated *ph*^*505*^ culture cells with the hormone, and quantified the expression level of several regions within the first intron at several developmental points. Compared to untreated *ph*^*505*^cells, the expression of both transcripts could be detected after exposure to ecdysone (Fig. 2B). One day after ecdysone treatment the expression of *let-B* transcript was the first to be observed, while *let-A* remained at basal levels (Fig. 2B). The expression level of *let-B* was similar for the first two days, and slightly increased in day three and four (Fig. 2B). On the other hand, the expression level of *let-A* was low during first two days after ecdysone treatment, but rapidly increased thereafter (Fig. 2B). In the tissue culture system induced by ecdysone it appears that the lncRNA *let-B*, containing all three EcREs, is the first to be expressed in response to ecdysone and *let-A* is expressed at a slightly later time point (see also Fig. 1A).

To confirm this sequential activation scheme *in vivo*, we analyzed how the two lncRNAs are expressed during fly metamorphosis. As the expression of the *let-7-C* locus is regulated by ecdysone signaling at the transition from larva to pupa, we collected wild-type larvae and pupae at different developmental time points, and quantified the expression level for different regions in the *let-7* intron (Fig. 2C). At late 3^rd^ larval instar (LL) and wandering larval (WL) stage, both of the two lncRNAs were expressed at a basal level, but started to increase expression at white pupae (WP) stage (Fig. 2C). Initially, the expression profiles of both lncRNAs follow the profile of other ecdysone induced genes like Eip75B. During the first 3 hours after pupae formation (APF) (P1-P3), the expression of *let-B* started again to decreased. In contrast, *let-A* expression continued to increase to a second peak at P3. The expression of both lncRNAs decreased at 24 hours APF, and we could only detect basal levels at 48 hours, 72 hours, and 96 hours APF (Fig. 2C). The expression dynamics of the two lncRNA shows significant differences to the expression pattern of the miRNA *let-7* (Fig. 2D). *In situ* hybridization of *let-7-C* first intron sequences to whole salivary glands of white prepupae show a strong signal at the periphery of the nuclei, reflecting most probably a highly active site of synthesis (Fig. 2E). These results show that the two lncRNAs are expressed transiently during early pupal stage in normal development and exhibit a different expression profile compared to that of the *let-7* miRNA.

### Effects of induced expression of the two lncRNAs in culture cells

To characterize the potential function of the two lncRNAs, we transduced *ph*^*505*^ culture cells with a silver-inducible viral vector expressing either *let-A* or *let-B*. In addition, we transduced either the mCherry sequence as control, or the entire second half of the first intron (named *I1B*) (Fig. 1B). In untransduced cells (Fig. 3A) or *ph*^*505*^ cell cultures treated only with silver (Fig. 3B), most cells had either an extended shape and attached to the plate (arrow in 3A), or a round shape and floated in the medium (arrowhead in 3A), with only few dead cells. However, when we induced the expression of *let-A*, surprisingly almost all the *ph*^*505*^ cells died within three hours (Fig. 3C, asterisks), with few remaining cells showing an abnormal shape (Fig. 3C, dashed arrow). When inducing *let-B, I1B*, or mCherry in *ph*^*505*^ cells, no such effects were observed (Fig. 3D-F). We used the alamarBlue assay to quantify cell viability after the different RNAs were induced. In contrast to the untreated control cells, cell viability was significantly reduced at three hours after *let-A* was expressed in *ph*^*505*^ cells (Fig. 3G). However, the cell viability was not affected when *let-B, I1B*, or mCherry was induced (Fig. 3G). To test if any of these RNAs might have a long-term effect, we measured the cell viability at three days after induction. As expected, most of the cells were dead when *let-A* was induced (Fig. 3H). While the expression of *let-B* only weakly reduced the cell viability, the induction of *I1B* or mCherry did not affect the cell viability (Fig. 3H). These results show that, unlike *let-B*, the expression of *let-A* is highly toxic to *ph*^*505*^ culture cells. Interestingly, the effect of *let-A* seemed to be cell type specific. When we transfected the RNAs in S2 or Cl8 cells and induced the expression, the cells did not die and the viability was not affected (Fig. 3I). As *let-A* showed such a strong phenotype to cultured cancer cells, we decided to focus on this lncRNA for further investigation.

**Figure 3.**
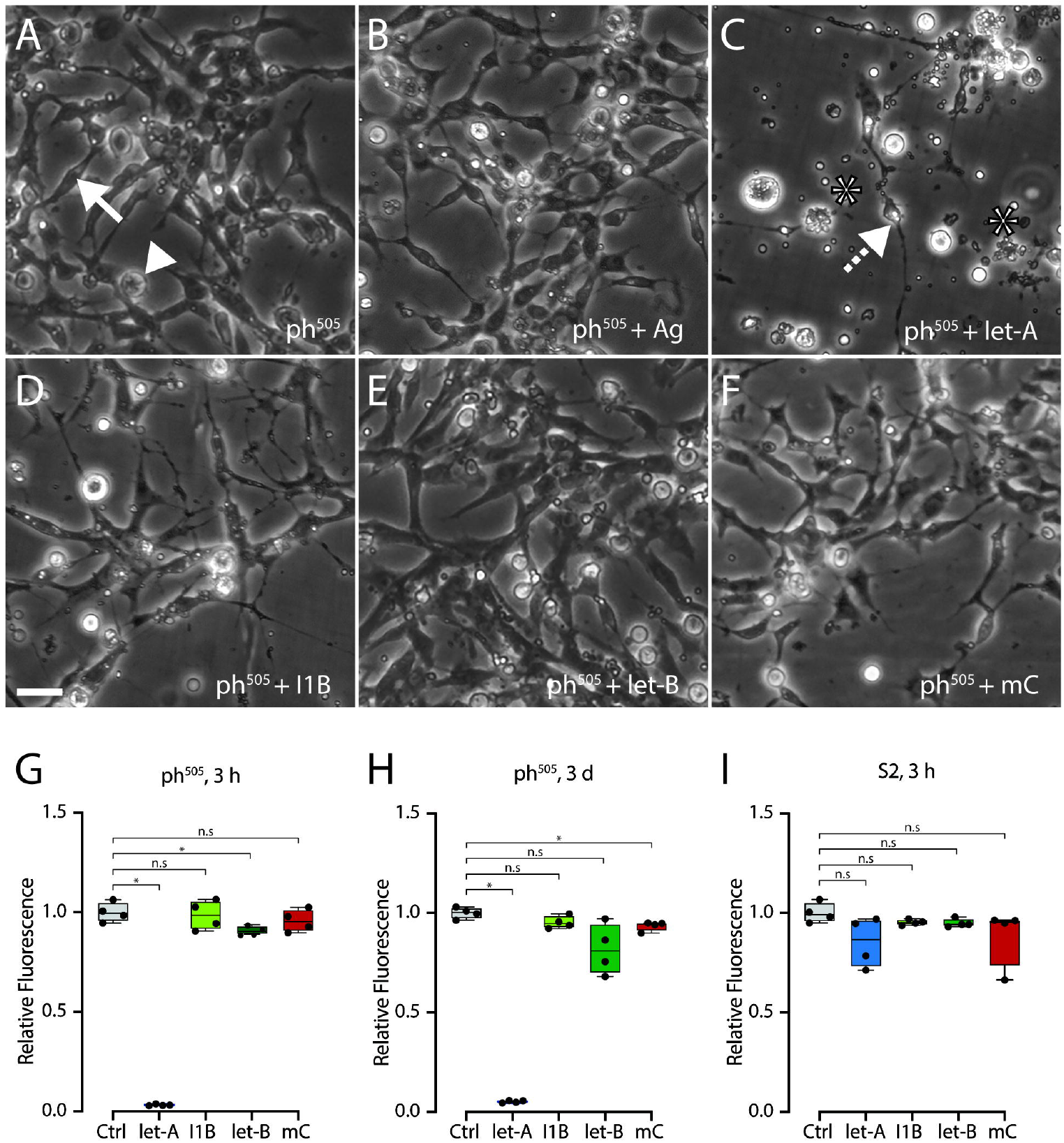
Induced expression of *let-A* resulted in rapid cell death in *ph*^*505*^ cells. (**A**-**F**) Light microscope pictures showing *ph*^*505*^ cells cultured in different conditions: (**A**) untransfected; (**B**) silver-treated; (**C**) induced *let-A*; (**D**) induced I1B; (**E**) induced *let-B*; (**F**) induced mCherry. All pictures were taken at two hours after induction. Arrow in (**A**) indicates a *ph*^*505*^ cell which has an extended shape and attaches to the plate; arrowhead in (**A**) indicates a round shape floating *ph*^*505*^ cell. Asterisks in (**C**) indicate dying cells; dashed arrow in (**C**) indicates a *ph*^*505*^ cell with abnormal shape. Scale bars is 20 μm. (**G**) alamarBlue quantification of cell viability at three hours after different RNAs were induced in *ph*^*505*^ cells. This assay incorporates an oxidation-reduction indicator that both fluoresces and changes color in response to chemical reduction of medium resulting from cell growth (Kumar et al., 2018). (**H**) alamarBlue quantification of cell viability at three days after different RNAs were induced in *ph*^*505*^ cells. (**I**) alamarBlue quantification of cell viability at three hours after different RNAs were induced in S2 cells. n.s. p ≥ 0.05, * p < 0.05 determined by Mann-Whitney test.

To determine how cells die upon *let-A* expression, we performed immunostaining for the apoptosis marker Annexin V. In the control culture, with mCherry expressed, only very few dead cells could be observed (Fig. 4A). In contrast, in *let-A* expressing cultures, as early as 40 minutes after induction, most of the cells were undergoing apoptotic cell death (Fig. 4B). Next, we tested if the cells die due to oxidative stress using the CellRox assay. Expression of mCherry did not induce any change in oxidative stress in the *ph*^*505*^ cells (Fig. 4C). However, when *let-A* was expressed in the *ph*^*505*^ cells, a significant increase in cellular oxidative stress could be observed (Fig. 4D). We next applied several apoptosis inhibitors against pan-caspases, caspase 8, Bax, or oxidative stress inhibitor to the *ph*^*505*^ cells during *let-A* expression. In all conditions, inhibition of apoptotic cell death resulted in more surviving cells in the culture (Fig. 4E). All together, these results show that induced expression of *let-A* in *ph*^*505*^ cells can lead to a rapid apoptotic cell death.

**Figure 4.**
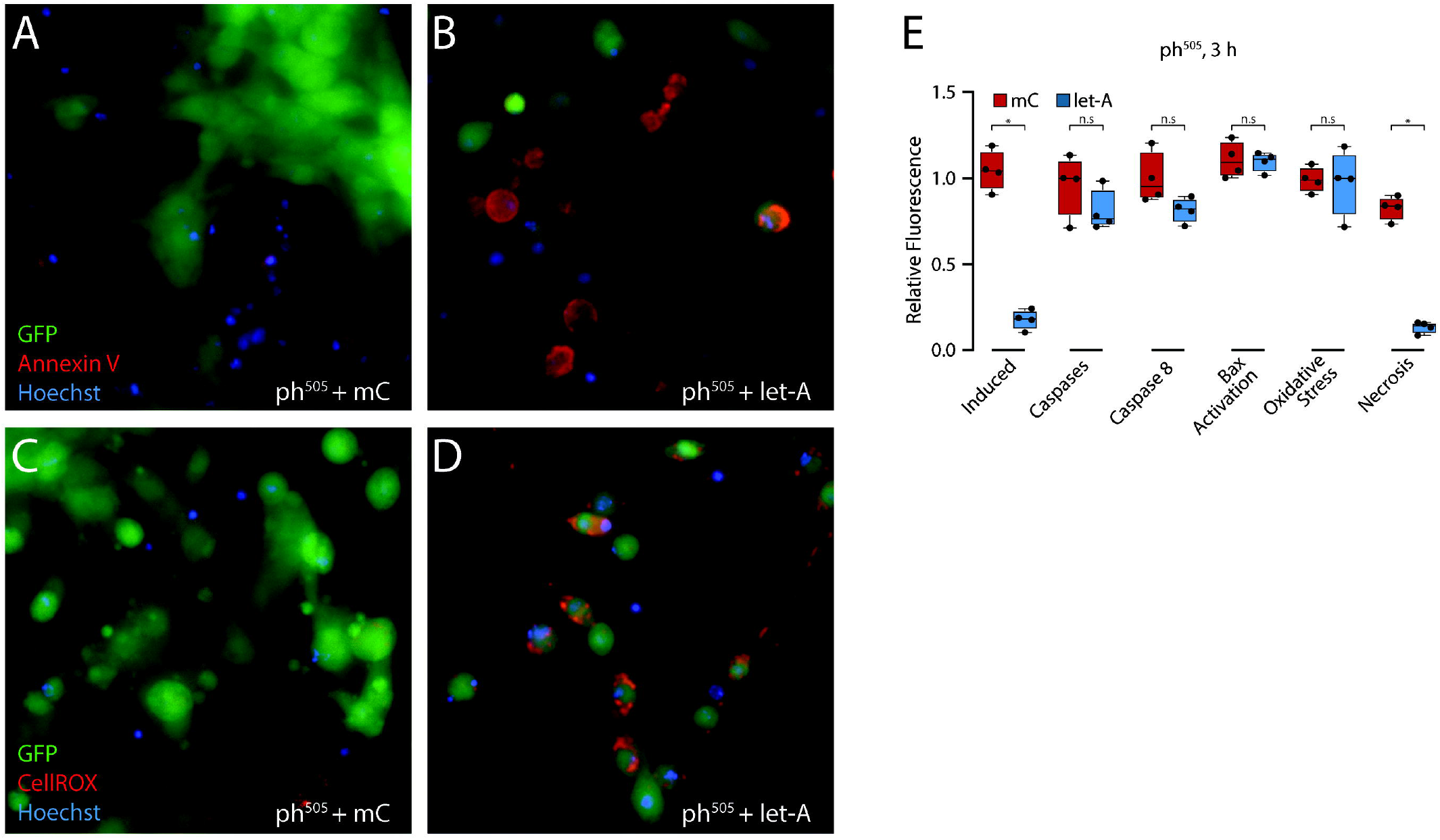
*let-A* expression induced apoptotic cell death in *ph*^*505*^ cells. (**A**) Annexin V staining in mCherry-expressing *ph*^*505*^ cells. (**B**) Annexin V staining in *let-A*-expressing *ph*^*505*^ cells showing many cells undergo apoptotic cell death. (**C**) CellRox staining in mCherry-expressing *ph*^*505*^ cells. (**D**) CellRox staining in *let-A*-expressing *ph*^*505*^ cells. (**E**) Cell viability measurements of *ph*^*505*^ cells when different apoptosis inhibitors were applied during the induction of mCherry (red) or *let-A* (blue). Scale bar is 10 μm. n.s. p ≥ 0.05, * p < 0.05 determined by Mann-Whitney test.

We have previously shown that *chinmo* is over-expressed in *ph*^*505*^ cells and acts as an important driver of tumorigenesis. Ecdysone signaling can down-regulate the expression of *chinmo* and transform tumorigenic *ph*^*505*^ cells into non-tumorigenic metamorphed cell. Similarly, when *let-A* was induced in *ph*^*505*^ cells, the expression of *chinmo* was also strongly down regulated (supplementary figure C). In contrast, induced expression of mCherry or *let-7-C* coding sequence did not reduce the expression of *chinmo* in *ph*^*505*^ cells. In addition, qPCR analysis of several other known *let-7* miRNA targets (*wech, cwo, Imp*) did not show down-regulation upon *let-A* induction (supplementary figure C).

### Cellular toxicity is connected to the *let-A* RNA

As only a small subset of *ph*^*505*^ cells was transduced by the viral vector (supplementary figure B), it was surprising to observe that almost all the cells in the culture died within a short time after *let-A* induction (Fig. 3C/G). We speculated that the transduced cells released some molecules, rendering the medium toxic and causing death in untransfected cells. To test this hypothesis, we induced expression of *let-A* (or mCherry RNA as control) in *ph*^*505*^ cells, collected the medium (henceforth called let-A/medium or mCherry/medium, respectively), and added the collected medium to another plate of untreated *ph*^*505*^ cells. Addition of the let-A/medium promoted death of most *ph*^*505*^ cells within the hour (Fig. 5A). Conversely, adding mCherry/medium had no effect (Fig. 5B). To test whether the medium toxicity was due to cell apoptosis in general, we killed untreated *ph*^*505*^ cells by UV irradiation, collected the UV/medium, and added it to a new plate of *ph*^*505*^ cells. Unlike the let-A/medium, the UV/medium had no immediate effect on *ph*^*505*^ cells (Fig. 5C), indicating the toxicity of the medium is due to cells with induced expression of *let-A*. To characterize the effector molecule in the medium, we induced the expression of *let-A* in *ph*^*505*^ cells and collected the medium after most cells had died. We next isolated either RNA or DNA from the medium by two rounds of purification and added the resulting fractions to *ph*^*505*^ cells. Purified RNA or DNA from the mCherry/medium did not show any effect to *ph*^*505*^ culture cells (Fig. 5D, E). In contrast, purified RNA from let-A/medium was able to kill all the cells (Fig. 5F), whereas DNA was not (Fig. 5G), suggesting that the toxicity was mediated by RNA molecules. To further confirm this result, we added RNase or DNase in the medium during the induction. Clearly, adding RNase to the medium reduced toxicity by restoring cell viability but DNase did not (Fig. 5H). Furthermore, cell viability tests showed that the toxicity was due to the specific expression of *let-A*, because purified RNA from the medium of other induced RNAs (*let-B, I1B*, or mCherry) could not kill *ph*^*505*^ cells (Fig. 5I). The toxicity of the RNA was also cell-specific similarly to the *let-A* transductions as there was no effect when S2 cells were treated with the purified RNA from *let-A* medium (Fig. 5J).

**Figure 5.**
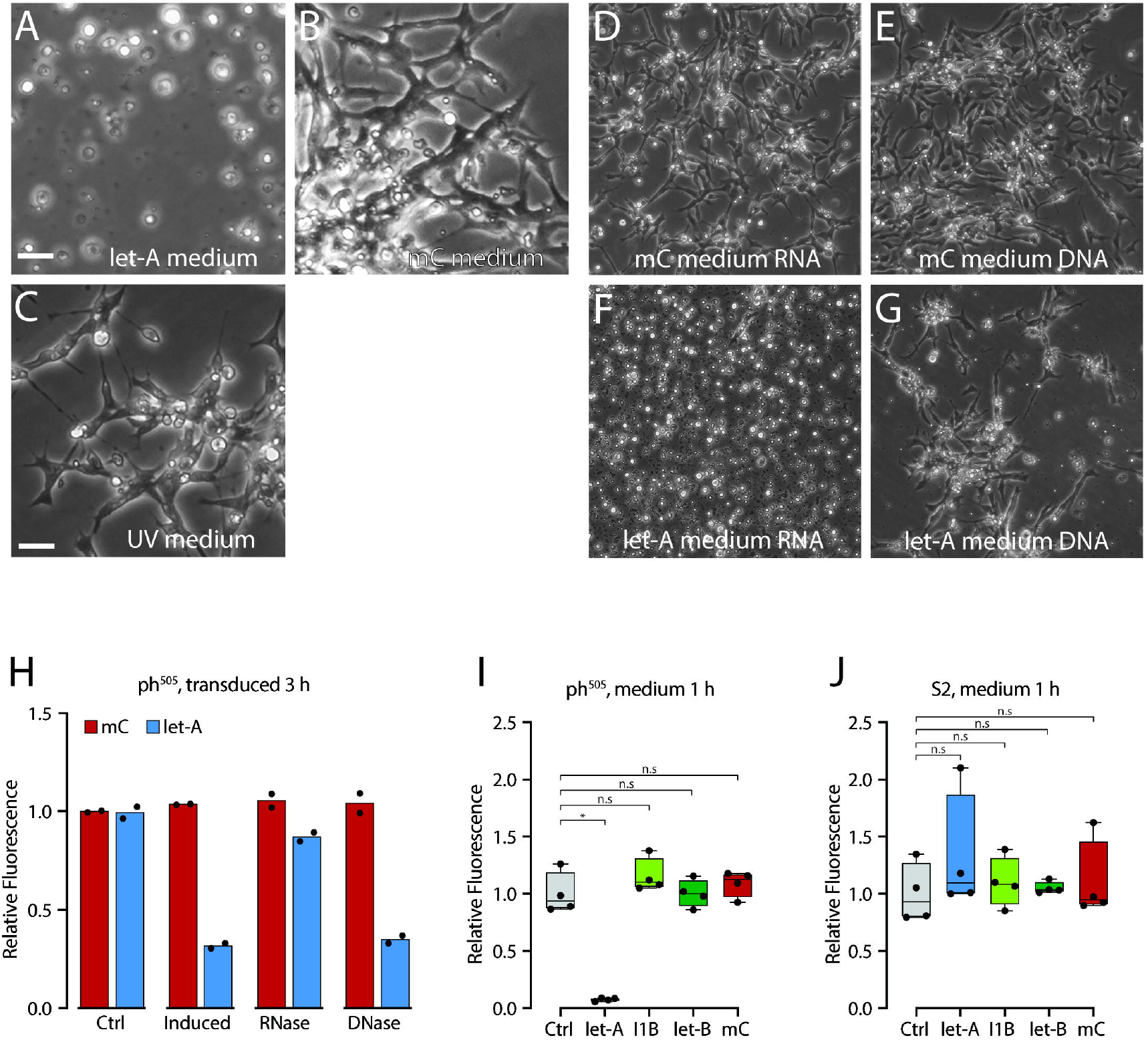
Cell death was induced by RNA molecules. (**A**-**C**) Light microscope images showing *ph*^*505*^ culture cells treated with let-A/medium (**A**), mCherry/medium (**B**), or UV medium (**C**). Scale bars are 20 μm. (**D**) Light microscope pictures showing *ph*^*505*^ culture cells treated with purified RNA fraction from mCherry/medium. (**E**) Light microscope pictures showing *ph*^*505*^ culture cells treated with purified DNA fraction from mCherry/medium. (**F**) Light microscope pictures showing *ph*^*505*^ culture cells treated with purified RNA fraction from let-A/medium. (**G**) Light microscope pictures showing *ph*^*505*^ culture cells treated with purified DNA fraction from let-A/medium. Scale bars are 40 μm. (**H**) Cell viability measurements of *ph*^*505*^ cells when RNase or DNase were added to the medium during the induction of *let-A* or mCherry. Note that adding RNase could significantly increase cell viability in the culture, but DNase could not. (**I**) Cell viability measurements of *ph*^*505*^ cells treated with RNAs purified from *let-A, let-B, I1B*, or mCherry/medium. Only purified RNA from let-A/medium was toxic to the cells. (**J**) Cell viability measurements of S2 cells treated with RNAs purified from *let-A, let-B, I1B*, or mCherry/medium. Purified RNA from let-A/medium was not toxic to S2 Cells. n.s. p ≥ 0.05, * p < 0.05 determined by Mann-Whitney test.

### let-A/medium and purified *let-A* RNA promote cell death of *ph*^*505*^ tumor tissue

Since *ph*^*505*^ culture cells can be killed by the *let-A*/medium and by purified RNA, we hypothesized whether the RNA may also be toxic to *in vivo* growing tumors. We first transplanted imaginal discs containing *ph*^*505*^ mutant clones into adult host flies and collected the tumors growing in those flies (Jiang et al., 2018; Torres et al., 2018). We dissected host flies carrying 1-week-old tumors and incubated the tumors with either mCherry/medium as control or let-A/medium for 24 hours. The tumor tissue changed morphology and became mildly dissociated after incubation with let-A/medium (Fig. 6A). In contrast, the cells in the tumor incubated in the control medium still formed tumor spheres and the bulk of tumor tissue did not show visible changes (Fig. 6B). Interestingly, after re-transplantation into adult host flies, let-A/medium-incubated tumors were no longer able to grow in host flies (n=20) (Fig. 6C), but the control medium-incubated tumors remained tumorigenic and gave rise to the formation of tumors in 75% of the hosts (n=20) (Fig. 6D). These results show that the let-A/medium is also toxic to *in vivo* formed tumors. Similarly, we found that tumors incubated with the let-A/medium-purified RNA became dissociated (Fig. 6E) and could not form tumors after re-transplantation. In contrast, tumors incubated overnight with RNA from control medium were still able to grow after re-transplantation (Fig. 6F).

**Figure 6.**
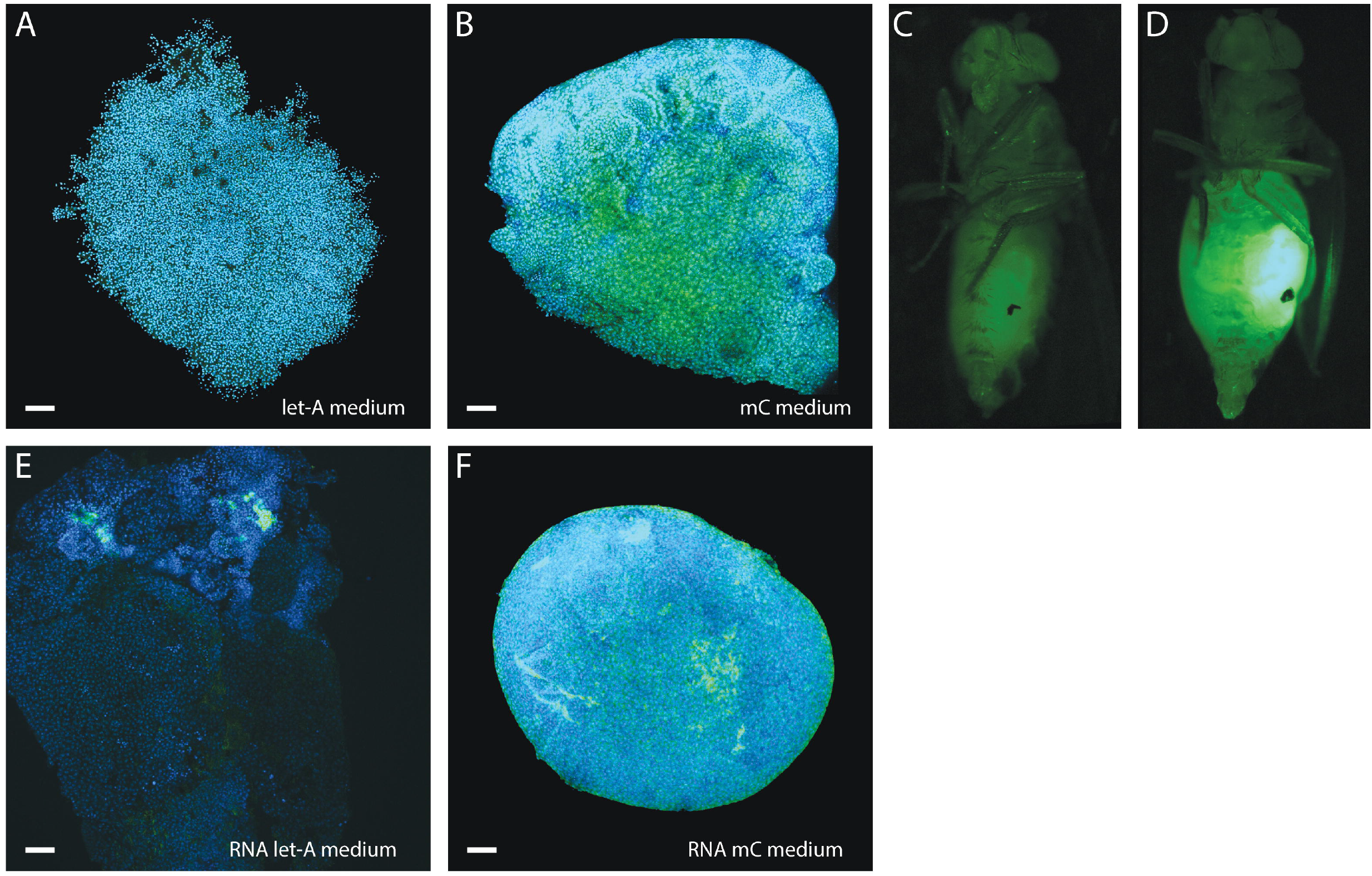
*let-A* was toxic to *ph*^*505*^ tumors grown *in vivo*. (**A, B**) Confocal images showing *ph*^*505*^ tumor after 24 hours incubation in let-A/medium (**A**) or mCherry/medium (**B**). Blue: DAPI staining; Green: GFP from *ph*^*505*^ cells. Note the tumor cells became dissociated and no longer formed a tight sphere structure after incubation in let-A/medium. Scale bars are 50 μm. (**C, D**) Adult host flies transplanted with *ph*^*505*^ tumor tissue incubated in let-A/medium (**C**) or mCherry/medium-incubated (**D**). let-A/medium-incubated *ph*^*505*^ tumor could not grow, but mCherry medium-incubated *ph*^*505*^ tissue could still retain tumor phenotype in the host flies. (**E, F**) Confocal picture showing *ph*^*505*^ tumor after overnight incubation with purified RNAs from let-A/medium (**E**) or from mCherry/medium (**F**). Note the tumor cells became dissociated and no longer formed a sphere structure after incubation with purified RNA from let-A/medium. Scale bars are 50 μm.

Next, we tested if the *let-A*/medium could directly affect the growth of tumors carried by flies. It is technically challenging to inject enough amount of medium or RNA into living flies. As an alternative, we performed a feeding assay using fly food prepared either with let-A/medium or mCherry/medium as control. Host flies carrying a small piece of tumor at one day after transplantation were then raised either on food with let-A/medium or with mCherry/medium. Flies were subsequently transferred to freshly collected medium-prepared food three times a week, and the growth of transplanted tumors in each fly was recorded during the entire period. Small pieces of tumor could grow and give rise to a large tumor mass after two weeks in all the hosts fed on food made with added mCherry/medium (Fig. 7A). However, the volume of the tumors in a subset of flies fed on let-A/medium food was visibly smaller (Fig. 7B). In about 1/3 of the host flies (n>50) fed with let-A/medium-prepared food, the growth of the tumor was slower and the volume of the tumor tissue was smaller (Fig. 7C, D). These results indicate that let-A/medium seems to have an effect on the growth of tumor tissue in a subset of flies.

**Figure 7.**
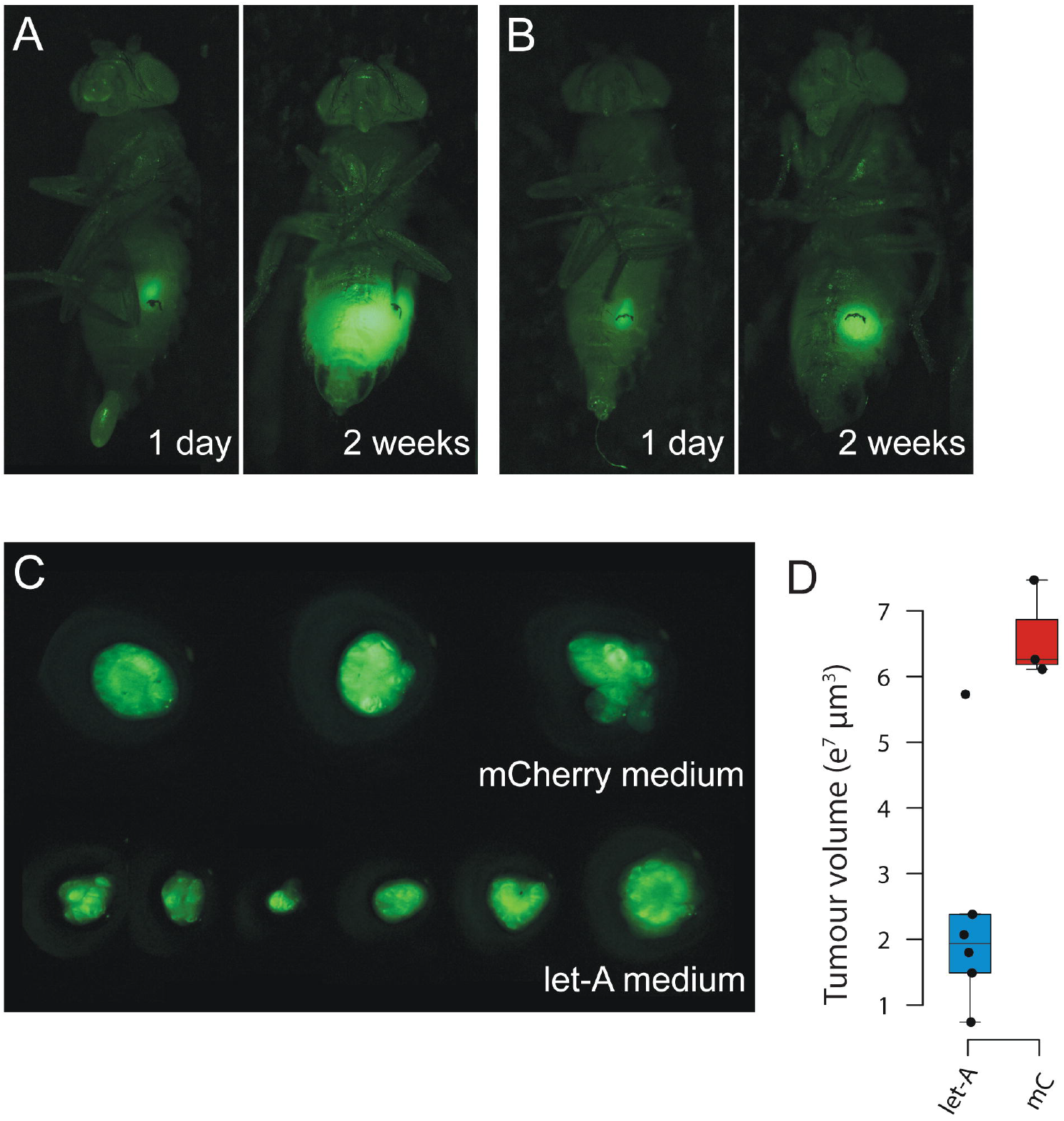
Feeding let-A/medium suppresses the growth of tumor in a subset of host flies. (**A**) Adult host flies transplanted with *ph*^*505*^ tumor (green fluorescent tissue in the abdomen) raised on mCherry/medium-containing food at one day or two weeks after transplantation. (**B**) Adult host flies transplanted with *ph*^*505*^ tumor (green fluorescent tissue in the abdomen) raised on let-A/medium-containing food at one day or two weeks after transplantation. (**C**) Dissected tumors from hosts raised either on mCherry/medium-containing food (upper row) or let-A/medium-containing food (lower row), note that 5 out of 6 tumors show reduced growth. (**D**) Quantification of the volume of the tumors shown in (**C**).

## Discussion

Most of the previous work on *Drosophila let-7-C* was focused on the role of the encoded miRNAs, in particular *let-7* (Chawla et al., 2016; Gendron & Pletcher, 2017; Sempere et al., 2002; Sokol et al., 2008). In this study, we show that in addition to the transcript encoding the three miRNAs, there are at least two lncRNAs expressed from the first intron. Our findings suggest the genomic organization, transcriptional regulation, and biological functions of the *let-7-C* to be much more complex than foreseen. Given that the clustering and arrangement of the encoded miRNAs is highly conserved the apparent novel functional units might in addition contribute to an evolutionary required cooperation between the various elements of the complex. We demonstrated previously that *let-7-C* plays an important role in a tumor cell eviction mechanism at metamorphosis (Jiang et al., 2018). We believe this to be a particular mechanism that evolved not specifically for eradicating cancer cells, but to eliminate undifferentiated cells at the transition between the larval and adult stage. Histolysis of larval structures and differentiation of imaginal cells to adult appendages is clearly a distinctive physiological event at metamorphosis, exemplifying this function. In the light that cancer cells often depict an undifferentiated state as a hallmark (de Thé, 2018; Torres et al., 2018), metamorphosis appears to be a bottleneck, leading to their specific destruction. The ecdysone-triggered activation of *let-7* production is a prerequisite for this process. *let-7* was shown in different organisms to act as a heterochronic gene, required to execute the transition from one developmental stage to the next (Bussing et al., 2008). Currently, we can only speculate what role the two newly identified lncRNAs may play in this mechanism. We attempted to identify the normal biological function of *let-A* by deleting corresponding parts of the locus using CRISPR/Cas9 technology. Unfortunately, all attempts to delete just *let-A* and leave the production of the other transcripts intact failed. We were not able to recover fly lines with the expected mutations. For the other lncRNA, *let-B*, we did not observe a similar effect when over-expressed in *ph*^*505*^ cells. Therefore, the true biological function of *let-B* remains unclear as well. We speculate, however, that the two lncRNAs play an important role in the eviction process of undifferentiated cells at the transition between the larval and adult stage and in consequence in the elimination of cancer cells we observed (Jiang et al., 2018). Both are activated by the ecdysone pulse. Interestingly, the *let-7-C* locus becomes clustered at the nuclear periphery in pupal salivary glands producing apparently large quantities of *let-A*. This suggests a presumptive role through secretion, possibly explaining the observed trans-effect on cancer cells. When *let-A* RNA was over-expressed in cultured *ph*^*505*^ cells, it resulted in rapid cell death of the entire population within a few hours. Similarly, when ph^505^ cells were treated with ecdysone, activating let-7-C expression, cells started to die, albeit at a much slower rate compared to a direct overexpression of *let-A*.

Induced expression of *let-A* can down-regulate the expression of *chinmo*, which acts as a strong driver of tumorigenesis. We have previously demonstrated that ecdysone signaling is fundamentally required for the process. The miRNAs (in particular *let-7* and *mir-125*) have an established role in cellular reprogramming, by for example inducing terminal cell cycle arrest (Caygill & Johnston, 2008). In consequence, *let-A* could be responsible for the apoptotic elimination of the undifferentiated cancer cell. We provided several lines of evidence that *let-A* was the molecule to induce the rapid and specific cell death. First, we thoroughly purified the different fractions from the let-A/medium and showed only the RNA fraction exerted toxicity. Second, RNase, unlike DNase, treated let-A/medium lost toxicity. Third, sonicating *let-A* RNA, or *in vitro* transcribed anti-sense RNA, abolished the toxic activity. Fourth, *in vitro* transcribed *let-A* RNA was sufficient to kill *ph*^*505*^ cells, although requiring a longer time and pre-incubation with cell extract (as we show in the accompanying paper (Birbaumer et al., 2021)). As many RNAs need post-transcriptional regulation to become active, it is plausible that *let-A* RNA needs further cellular modifications and/or processing.

As *let-A* RNA shows a strong toxic effect on cultured *Drosophila* tumor cells, we were attempted to further test its application on *in vivo* grown tumors. Injecting RNA into living flies is technically challenging, and the wounds generated during injections reduce viability. By alternative means, we first show tumors become non-tumorigenic after incubated with let-A/medium, as they can no longer grow in size after re-transplantation into hosts. In addition, feeding flies with let-A/medium leads to reduced growth of the transplanted tumors in a subset of hosts. It is surprising that the RNA appears to be stable in the feeding tests, requiring additional research efforts to understand the reason for the stability and the mechanism how the *let-A* RNA acts in organism. In consequence, we were wondering whether the oncolytic activity of *let-A* is evolutionary conserved. Indeed, in our accompanying paper (see (Birbaumer et al., 2021)), we demonstrate *let-A* to have an oncolytic effect on various types of human cancer cells. These results therefore open the possibility to consider this molecule for a novel tumor therapeutic treatment.

## Supplementary Figure

A. Overexposing of the Northern blot in Fig. 1C, showing additional shorter bands that might be isoforms.

B. Transducing efficiency of the viral vector, showing a small subset of *ph*^*505*^ cells was transduced by the vector.

C. qPCR analyses showing the expression of *chinmo* was also strongly down regulated after let-A expression. In contrast, induced expression of mCherry or *let-7-C* pri-RNA coding sequence did not reduce the expression of *chinmo* in *ph*^*505*^ cells. In addition, several other known *let-7* miRNA targets (*wech, cwo, Imp*) did not show down-regulation upon *let-A* induction.

## Materials and Methods

### Fly genetics

Fly strains were maintained on standard medium. All genetic experiments were performed at 25°C. To generate homozygous *ph*^*505*^ MARCM clones, virgin females of *ph*^*505*^ *FRT19A/FM7 act-GFP* were crossed with *tub-Gal80 FRT19A; ey-flp act>STOP>Gal4 UAS-GFP* males.

To measure the expression of the lncRNA during development, we collected larvae and pupae (*w*^*1118*^) at different developmental time points and extracted RNA for qPCR analyses. For larvae, third instar larvae staying in the food were collected as late larvae (LL); and wandering larvae (WL) were larvae leaving the food and wandering in the vial. For pupae, white pupae (WP) were collected and aged in a separate plate for 1, 2, 3, 24, 48, 72, and 96 hours.

The following fly strains were used in this study: (1) *Ore-R*, (2) *w*^*1118*^, (3) *ph*^*505*^ *FRT19A/FM7 act-GFP*, (4) *tub-Gal80 FRT19A; ey-flp act>STOP>Gal4 UAS-GFP*.

### Tissue transplantations

Transplantation experiments were carried out as previously described (Jiang et al., 2018). In brief, 4–6 days old adult *w*^*1118*^ females were used as hosts (n>20 for each transplantation). The host flies were immobilized on an ice-cold metal plate and stuck on a piece of double-sided sticky tape, with their ventral sides up. The dissected tumor tissue was cut into small pieces of similar size and each piece was transplanted into the abdomen of one host using a custom-made glass needle. All transplantation was made under a GFP microscope to ensure labeled cells were injected into the hosts. After transplantation, host flies were allowed to recover at room temperature for 1-2 hours in fresh standard *Drosophila* medium before transferred to and maintained at 25°C.

To test the toxicity of the let-A/medium on tumors grown in the abdomen, host flies carrying 1-week-old transplanted *ph*^*505*^ tumor tissue were dissected, the tumors removed and incubated with either control medium (mCherry/medium) or let-A/medium for 24 hours. The tumors were then cut into small pieces and re-transplanted into new host flies.

### *ph*^*505*^ cell line generation

To establish *ph*^*505*^ cell line, two to three 1 week-old transplanted *ph*^*505*^ tumors were isolated from the host flies and dissected into small pieces in sterilized PBS following 3 times wash with Shields and Sang M3 Insect medium (US Biological) containing Penicillin/streptomycin(P/S). The cells were resuspended in 1ml of M3 (P/S) medium and plated into the 96 well plate 200µl per well. After stable cell growth, a single cell line was isolated using Terasaki plate (Greiner Bio-One). To confirm the identity of the cell line, genomic DNA was isolated and sequence analysis was performed.

### RNA *in situ* in salivary glands

For single molecule FISH analysis (Fluorescence *in situ* Hybridization) a set of oligonucleotides spanning the first intron of *let-7-C* was designed using the Stellaris Probe Designer software (Biosearch Technologies). The oligos were ordered from Biosearch Technologies labeled with Quasar-670.

RNA FISH was preformed following the Stellaris protocol for suspension cells with minor modifications. Briefly, salivary glands from white pupa were dissected and washed in PBS and fixed with 4% formaldehyde in PBS for 10 min at room temperature. After washing with PBS samples were incubated with 70% EtOH for 24 hours at 4°C. Then, samples were washed in washing buffer (2xSSC, 10% formamide) at room temperature. For hybridization 125 nM of the probe set in hybridization buffer (Biosearch Technologies) was added and incubated overnight at 37 ^0^C. The next day, samples were washed twice for 30 min at 37 ^0^C with washing buffer, the second wash containing 500 ng/ml DAPI (Sigma□Aldrich). After one wash with PBS samples were mounted with Vectashield Mounting Medium (Vectorlabs).

### Cell culture, treatments and viability

*Drosophila* Clone 8 (Cl.8) and *ph*^*505*^ cell lines were maintained at 25 °C in Shields and Sang M3 Insect medium (US Biological), supplemented with 2% Fetal Bovine Serum (FBS) (PAN Biotech), 2.5% fly extract, 5□µg/ml human insulin (Sigma), 100 U/ml penicillin and 100□μg/ml streptomycin (Gibco, Life Technologies). S2 cells were cultured in Schneider’s medium (Gibco, Life Technologies) supplemented with 10% FBS (PAN Biotech) at 25 °C. *Spodoptera frugiperda* Sf21 cells (Clontech) were propagated in Grace’s medium (Gibco, Life Technologies) with 10% FBS (PAN Biotech) at 27°C. SEAP activity was detected using QuantiBlue (InvivoGen) and the absorbance quantified on a Tecan Infinite M1000 PRO microplate reader at 650 nm. For the ecdysone time course, cells were treated one day after seeding with 5 μM 20-hydroxy-ecdysone (Sigma) and incubated for the specified time before RNA was extracted.

To induce apoptosis, *ph*^*505*^ cells were treated twice with UV-C (90 mJ/cm2, 254 nm). For the UV treatment, cells were washed in PBS and irradiated with UV light in PBS, which was afterwards replaced by medium. For the RNase and DNase treatment 100 μg/ml RNase mix (Roche) or 100 U/ml DNase I (Roche) and 100 μM AgNO3 were added together to the transduced cells. After 3 hours cell viability was measured using alamarBlue.

The following inhibitors were used: Apoptosis inhibitors: Pan-Caspase Inhibitor Z-VAD-FMK (Enzo Life Sciences) 50 μM; Caspase 8 Inhibitor Z-IETD-FMK (Enzo Life Sciences) 15 μM; Clusterin (secretory form, human recombinant, Enzo Life Sciences) 0.5 μM; oxidative stress was reduced with 1 mM Trolox (6-Hydroxy-2,5,7,8-tetramethylchromane-2-carboxylic acid, Sigma). For all the inhibitor experiments, cells were pretreated one hour with the respective inhibitor before induction or addition of the extracted medium. For induction, the cell viability was determined after three hours, for extracted medium treatment after one hour. Cell viability was measured after the indicated time using alamarBlue cell viability assay reagent (Thermo Fisher Scientific) following the manufacturer’s instructions. Fluorescence at 650 nm was measured with Tecan Infinite M1000 PRO microplate reader.

### Generation of recombinant baculoviruses and transduction of *Drosophila* cells

Cloning was performed following the strategy described in (Lee et al., 2000). The *let-A* and *I1B* fragments were amplified from S2 cell genomic DNA using the Expand Long Template PCR System (Roche) and cloned in pBacPAK8_EGFP under a metallothionein promoter. *let-B* was cloned from pBacPAK8_EGFP_I1B. Viruses were generated, amplified, analyzed and harvested using the BacPAK Baculovirus Expression System (Clonetech) according to manufacturer’s instruction.

Cells were plated one day prior to transduction. For transductions cells were first washed twice with Grace’s medium and incubated for two hours at room temperature with the virus inoculum. After removal of the virus, fresh medium was added and cells were incubated at 25°C for one more day before induction. Unless otherwise specified, transcription of the constructs was induced with 100 μM AgNO3 (Sigma). The conditioned medium was always harvested one day after induction.

### Northern blot analysis

PolyA^+^ RNA was isolated using NEBNext Poly(A) mRNA Magnetic Isolation Module (NEB) and 1 ug total RNA or 0.1 ug polyA+ RNA were run on a 0.8% formaldehyde-agarose gel. Northern blotting was performed using DIG Northern Starter Kit (Roche) following manufacturers instructions.

### RNA extraction, mapping, cDNA synthesis and Quantitative real-time PCR

RNA from cells or medium was extracted using TRIzol (Invitrogen) following manufacturer’s instructions. RACE experiments were carried out using the SMARTer RACE 5’/3’ Kit (Clonetech) according to manufacturers instructions. As template total and poly A+ RNA from mixed *Drosophila* pupal stages was used.

Reverse transcription for qPCR analysis was done with the first strand synthesis kit (Fermentas) using oligo (dT)18 as a primer and qPCR was carried out with LightCycler 96 (Roche) using FastStart Essential DNA Green Master Mix (Roche). Expression levels were normalized to ATPasecf6.

For miRNA expression analysis, RNA was reverse transcribed using TaqMan miRNA Reverse Transcription kit (Applied Biosystems) and qPCR was performed using let-7 and 2S rRNA TaqMan MicroRNA Assay primer mix (Thermo Fisher Scientific) with the TaqMan Small RNA Assay (Applied Biosystems).

### Annexin V and CellROX staining

To detect an early stage of apoptosis, cells were incubated one day after transduction with 2.5 mM CaCl2 (Sigma), 5 μl/ml Alexa Fluor-conjugated Annexin V (Biolegend) and 1 μg/ml Hoechst33342 for 10 minutes at room temperature and then induced with 100 μM AgNO3 and imaged on several time points. To detect the presence of ROS, CellROX DeepRed Reagent (Life Technologies) was used. Before induction, cells were incubated for 30 minutes with 2.5 μM CellROX reagent. Nuclei were also stained with 1 μg/ml Hoechst33342. Images were taken using a Leica MZ16 FA fluorescent microscope.

### Immunohistochemistry and antibodies

Tumors were fixed in 2% paraformaldehyde (in 1xPBS) for 25 minutes at room temperature, and washed several times in PBST (1xPBS with 0.5% Triton X-100). Tumors were incubated overnight with primary antibodies at 4°C, followed by several washes at room temperature, incubated with secondary antibodies at 4°C overnight. After several washes, samples were incubated with DAPI (1:200 in PBST) at room temperature for 20 minutes, then mounted in Vectashield and stored at -20°C before imaging.

Primary antibody used in this study was: chicken anti-green fluorescent protein (GFP) (1:1000; Abcam ab13970); Secondary antibodies were: Alexa 488-conjugated anti-chicken (1:500; Molecular Probes).

Immunofluorescent images were recorded on a Leica TCS SP8 confocal microscope. Adult flies with transplanted tumors were taken on a Nikon SMZ1270 microscope. Images were processed using ImageJ, Imaris, Photoshop, and Adobe Illustrator.

### Transcriptome analyses

Transcriptome analyses were performed by mRNA sequencing. RNAs were isolated from the ph505 culture cells at 16 hours after treated with ecdysone (5 μM 20-Hydroxyecdysone (Sigma)). RNA was extracted using an Arcturus PicoPure RNA Isolation kit (Applied Biosystems), library prepared with SMARTSeq2 NexteraXT and sequenced on an Illumina NextSeq500.

### Bioinformatics analyses

Short reads were aligned to BDGP dm6 genome assembly using TopHat 2.0.12 for Bowtie 2.2.3 with parameters “--very-sensitive”. Conversion to bigWig was performed using R (3.5.1). RPKM values for external miRNA expression data were retrieved from FlyBase (Release 2018_06).

### Data availability

Deep sequencing data for metamorphed and eye disc cells can be found in GEO with accession GSE101455 and for time course data with accession GSE186539.

## Acknowledgments

This research was supported by ETH Zürich. We thank Martin Fussenegger for hosting TB, YJ and RP in his laboratory and Andreas Hierholzer for comments on the manuscript. Sequencing was performed by the Genomics Facility Basel.

