## Supplementary Figure for "A long non-coding RNA in the *let-7* complex acting as a potent and specific death effector of cancer cells": Supplementary Figure.pdf

A. Overexposing of the Northern blot in Fig. 1C, showing additional shorter bands that might be isoforms.

B. Transducing efficiency of the viral vector, showing a small subset of $ph^{505}$ cells was transduced by the vector.

C. qPCR analyses showing the expression of *chinmo* was also strongly down regulated after let-A expression. In contrast, induced expression of mCherry or *let-7-C* pri-RNA coding sequence did not reduce the expression of *chinmo* in $ph^{505}$ cells. In addition, several other known *let-7* miRNA targets (*wech*, *cwo*, *Imp*) did not show down-regulation upon *let-A* induction.

### Supplementary Figures

A

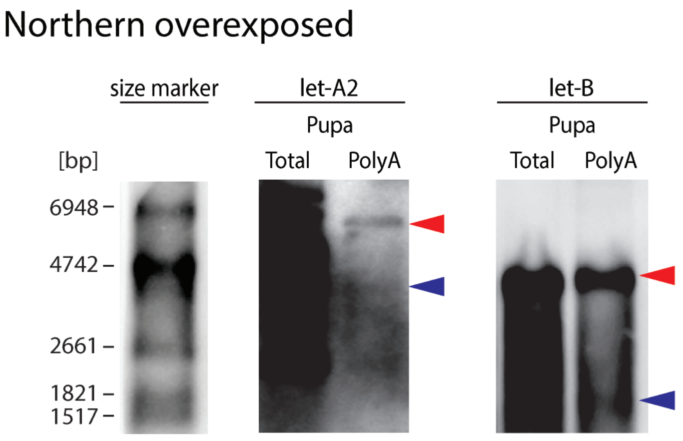


B

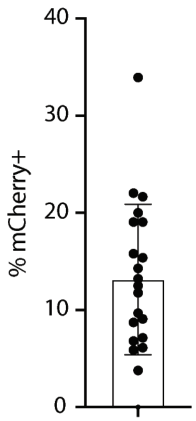


C

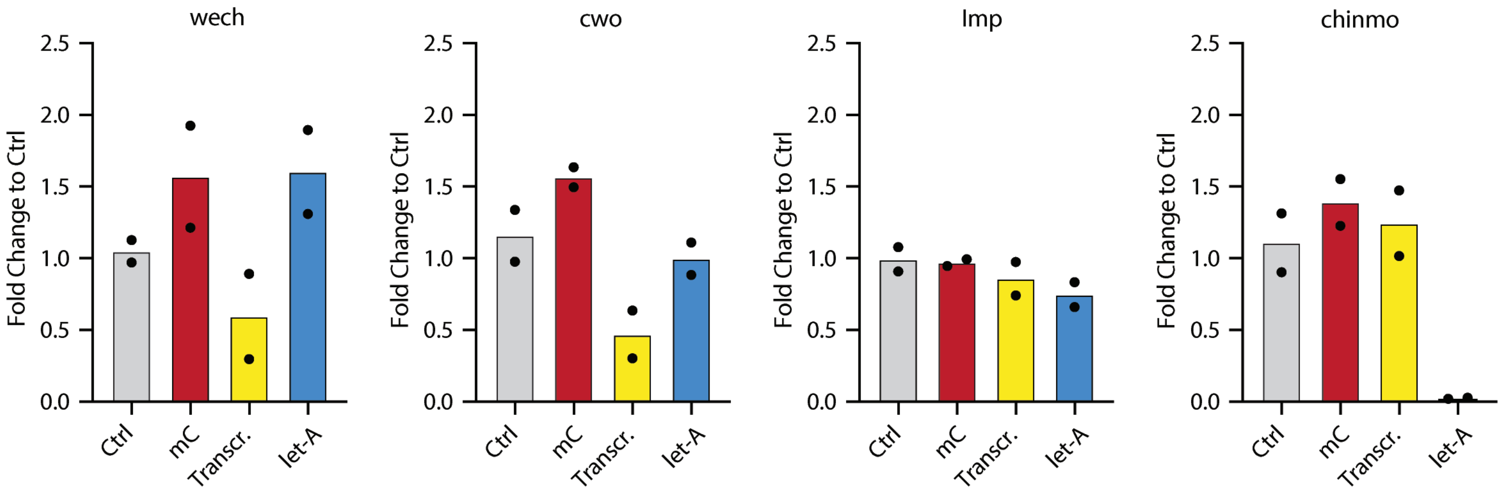
